# A Pan-Coronavirus Vaccine Candidate: Nine Amino Acid Substitutions in the ORF1ab Gene Attenuate 99% of 365 Unique Coronaviruses: A Comparative Effectiveness Research Study

**DOI:** 10.1101/2022.04.28.489618

**Authors:** Eric Luellen

## Abstract

**Background:** The COVID-19 pandemic has been a watershed event. Industry and governments have reacted, investing over US$105 billion in vaccine research.^1^ The ‘Holy Grail’ is a universal, pan-coronavirus, vaccine to protect humankind from future SARS-CoV-2 variants and the thousands of similar coronaviruses with pandemic potential.^2^ This paper proposes a new vaccine candidate that appears to attenuate the SARS-Cov-2 coronavirus variants to render it safe to use as a vaccine. Moreover, these results indicate it may be efficacious against 99% of 365 coronaviruses. This research model is wet-dry-wet; it originated in genomic sequencing laboratories, evolved to computational modeling, and the candidate result now require validation back in a wet lab.

**Objectives:** This study’s purpose was to test the hypothesis that machine learning applied to sequenced coronaviruses’ genomes could identify which amino acid substitutions likely attenuate the viruses to produce a safe and effective pan-coronavirus vaccine candidate. This candidate is now eligible to be pre-clinically then clinically tested and proven. If validated, it would constitute a traditional attenuated virus vaccine to protect against hundreds of coronaviruses, including the many future variants of SARS-CoV-2 predicted from continuously recombining in unvaccinated populations and spreading by modern mass travel.

**Methods:** Using machine learning, this was an *in silico* comparative effectiveness research study on trinucleotide functions in nonstructural proteins of 365 novel coronavirus genomes. Sequences of 7,097 codons in the ORF1ab gene were collected from 65 global locations infecting 68 species and reported to the US National Institute of Health. The data were proprietarily transformed twice to enable machine learning ingestion, mapping, and interpretation. The set of 2,590,405 data points was randomly divided into three cohorts: 255 (70%) observations for training; and two cohorts of 55 (15%) observations each for testing. Machine learning models were trained in the statistical programming language R and compared to identify which mixture of the 7.097 × 10^23^ possible amino-acid-location combinations would attenuate SARS-CoV-2 and other coronaviruses that have infected humans.

**Results:** Contests of machine-learning algorithms identified nine amino-acid point substitutions in the ORF1ab gene that likely attenuate 98.98% of 365 (361) novel coronaviruses. Notably, seven substitutions are for the amino acid alanine. Most of the locations (5 of 9) are in nonstructural proteins (NSPs) 2 and 3. The substitutions are alanine to (1) valine at codon 4273; (2) leucine at codon 5077; (3) phenylalanine at codon 2001; (4) leucine at codon 372; (5) proline at codon 354; (6) phenylalanine at codon 2811; (7) phenylalanine at codon 4703; (8) leucine to serine at codon 2333; and, (9) threonine to alanine at codon 5131.

**Conclusions:** The primary outcome is a new, highly promising, pan-coronavirus vaccine candidate based on nine amino-acid substitutions in the ORF1ab gene. The secondary outcome was evidence that sequences of wet-dry lab collaborations – here machine learning analysis of viral genomes informing codon functions -- may discover new broader and more stable vaccines candidates more quickly and inexpensively than traditional methods.

## Introduction

### Background

As of February 2022, the COVID-19 pandemic has killed at least 5.82 million people.^3^ Another 412 million people have been sickened, of whom an estimated 124 million (30%) will likely suffer long-term vascular illnesses leading to premature deaths.^3 4^ And, both numbers will continue to climb until an open-access or open-source vaccine is globally available.

While new to living generations, viral pandemics have repeatedly been mass casualty events in the history of humankind. Since 1900, viral pandemics have killed an estimated 187 million people.^5^ Medical historians believe viral pandemics have killed as many as 500 million people in the last 2000 years.^5^ Moreover, the risk of future viral pandemics is high and increasing, with five new infectious diseases emerging annually, each with pandemic potential.^6^ “There’s a huge inventory of potential [new viruses] …lurking in bats and civets and other animals; it’s only a matter of time before another one of them jumps into us.”^7^

> *“A growing body of scientific evidence, considered together with ecological reality, strongly suggests that novel coronaviruses will continue to infect bats and other animal reservoirs and potentially emerge to pose a pandemic threat to humans. To counter future coronavirus outbreaks, the global scientific and medical research community should focus a major effort now on…the development of long-lasting, broadly protective coronavirus vaccines*.*”*^8^

In response, global organizations are raising $3.5 billion, including $300 million from the Gates Foundation, to learn to develop new vaccines in an open-access format as quickly as possible.^9^ The Gates Foundation has donated $200 million to the search for such a universal, or pan-coronavirus, vaccine [AUTHOR’S NOTE: The Gates Foundation was sent a pre-print of this paper before Gate’s statement].^10^

This study was born of capability, curiosity, and this urgent necessity. The author’s training and prior research applying machine learning to identify coronavirus candidate immunotherapies informed the question of whether artificial intelligence could also categorize and prioritize which coronaviruses have zoonotic – species jumping – pandemic potential.^11^ Therefore, he sought to develop a pandemic early-warning system using machine learning that could autonomously screen biobanks for newly discovered coronaviruses that could infect humans and alert them when it found them to buy time to predict, prevent, or prepare for future pandemics. He succeeded to a high 90^th^ percentile level of accuracy and shared that information with interested national security officials in the United States in the first half of 2021. In the second quarter of 2022, he reverse-engineered this machine learning algorithm to learn which amino acids and genomic site substitutions likely attenuate SARS-CoV-2, then other coronaviruses, to design a new, broader, attenuated vaccine candidate for coronaviruses.

Recently, experts’ calls to action to the likelihood that the COVID-19 pandemic will continue for years has increased.^12^ The root cause is because poorer countries lack access to patented, higher-cost gene therapies, tests, and vaccines. The global urgency for open-access, open-science, and open-source coronavirus prevention solutions has reached such a tenor that when a team of Texan scientists recently identified a 90% effective vaccine against just the coronavirus variants that cause COVID-19 – approximately 1% of coronaviruses with pandemic potential – and donated the intellectual property, their Congresswoman nominated them for the 2022 Nobel Peace Prize. While the formula has won emergency approval in India, which has bought 300 million doses for $2 each from Indian pharmaceutical manufacturers, its efficacy data has yet to be accepted, reviewed, and published in a peer-reviewed scientific journal.^13^ That solution cost an estimated $7 million and two years to develop.^13^

By comparison, this paper intends to report in an open-access peer-reviewed scientific journal the results and formula for a vaccine candidate that addresses approximately two orders of magnitude more coronaviruses with 9% greater efficacy. It was developed over four months – three months in early 2021 and one month in 2022 – a time-value cost of approximately $100,000. While patent-pending, the patent will be waived for development by any country, not only low-or-middle income countries.

To maximize transparency, it is necessary to note that these results were initially offered to one American pharmaceutical company to license without disclosing the details here; however, during the slowness of their response, the author reconsidered the ethics of the private-vaccines-for-profit model versus global vaccine access during a pandemic. Altruism won. The author intends to offer this design in an open-access publication in an attempted effort to defeat or reduce vaccine nationalism, which could cost the global economy up to $1.2 trillion per year in the gross domestic product (GDP).^14^

The viral family of coronaviruses was chosen as the focus of this research for two reasons. One, their investigation has become urgent since the COVID-19 pandemic began. Two, the diversity of unidentified coronaviruses with zoonotic pandemic potential likely exceeds 3,204, excluding variants. Therefore, coronaviruses are a likely source of future pandemics.^15^

Non-structural proteins (NSPs) were chosen as a sub-focus because of five reasons: (1) they are more conserved and thus enduring and stable, making them less of a moving target for vaccines when compared to the ever-evolving spike (S) and membrane (M) genes; (2) they constitute approximately 67% of coronaviruses’ genetic material; (3) NSPs direct the RNA synthesis and processing in coronaviruses – they are at the root, or origin, of encoding and programming downstream pathogenesis; (4) research into other viruses has found attenuation solutions in the NSPs (e.g., Avian coronavirus, Chikungunya, Zika); and, (5) comprehensive reviews of coronavirus mutations have found that meaningful recombination results in variants rarely occur in the ORF1ab gene, making NSP attenuation less susceptible to evasion.^16 17 18 19^ Therefore, the ORF1ab gene and polyprotein are relatively more present and consistent across coronaviruses and time, decreasing the probability of vaccine evasion.^20 21^

A high-level understanding of coronaviruses’ NSPs, the ORF1ab gene, and polyprotein targeted here, will be helpful. ORF1ab expresses two polyproteins – ORF1a and ORF1b – that proteolytically cleave to form 11-16 NSPs, depending on the coronavirus. Generally, ORF1a drives the production of proteins involved in the early control of hosts’ innate immune response at the cellular level. ORF1b produces proteins to replicate the genome and synthesize RNA.^22^

### Goals of this Study

The goals of this study were to learn three things: (1) can a model be developed to predict which coronaviruses can infect humans and, if so, at what level of accuracy; (2) therein, the relative import of each codon, or variable, in making such predictions; and, (3) specifically, which amino acids that are encoded by those codon-locations can be substituted with what other amino acid to attenuate the virus’ ability to infect humans.

Investigating the etiology of a disease is essential in discovering new and more efficacious treatments. The premises behind this paper are that: (1) to make pathogens less dangerous, we must first understand the functional proteomics as to what makes them dangerous – to identify codons by their functionality; (2) pathogens’ dangers stem from their ability to infect and virulence; (3) these insights are prerequisites to precision genetic engineering reaching its potential benevolence; (4) it could be more economical and effective to combat infectious diseases and optimize the beneficence of viruses by editing their genes instead of editing hosts’ genes, which is more complicated, controversial and potentially dangerous; and, (5) a clinically accurate and scalable codon targeting system is a prerequisite for the most durable, efficacious and efficient attenuations or gene edits to enable viral vaccines (as opposed to Moderna and Pfizer’s preventative gene therapies, often mischaracterized as vaccines).

This study hypothesized that the advents and confluence of genomic sequencing, gene editing, and machine learning in moist labs – a collaboration of dry and wet labs – could precisely identify the functional locations, or codons, where proteins are synthesized in viruses to enable the infection of specific species and which amino acid need to be substituted to attenuate them. Here, that species is humans; a modification of the method could create a new model for any species of interest (e.g., crops, livestock, etc.). In other words, the end goal was to identify the upstream causes of human virulence in coronaviruses and how to attenuate them, ideally in such a generalized method that it may work for other viruses too.

## Methods

This comparative genomics study used computational biology, specifically, machine learning. This study was based on secondary data published by the NIH’s National Library of Medicine (NLM) and the National Center for Biotechnology Information’s (NCBI) database, Nucleotide, which amalgamates genomic sequences from GenBank, RefSeq, Third-party Annotation (TPA), and Protein Data Bank (PDB) databases.^23^ Therein, nucleotides were sequenced in 365 novel coronaviruses, of which 167 infected humans and 198 infected other species, including bats, birds, camels, civets, cows, and pigs.

First, the data set was pre-processed in three ways: (1) the codon values were transformed via a proprietary method to improve their digestibility by machine-learning algorithms; (2) each codon’s encoding was transformed to enable interpretation of the amino acid; and (3) coronavirus strains that were sampled from animals were labeled “0,” and coronavirus strains that were sampled from humans, evidencing their ability to infect humans, were labeled “1.” Each observation was also manually labeled with the scientific and common names of the source species, the number of base pairs in the gene, and the geographic location of the collection to ensure diversity within the dataset and minimize data bias.

Second, three types of primitive machine-learning algorithms – a kind of artificial intelligence based on brute-force statistics – and a machine-learning ensemble were applied to the data set of 365 observations of approximately 7,097 codons, or trinucleotides, of unique coronavirus genomic sequences of a single gene (ORF1ab). The Rattle library, version 5.3.0 (Togaware), in the statistical programming language R, version 3.6.3 x64 (CRAN), was used to apply these four types of machine-learning algorithms – an extreme gradient boosting (XGBoost), random forests of classification trees, nonlinear support vector machines (SVM), and deep-learning neural networks – to learn which algorithm, if any, could build a highly accurate predictive model. Rattle randomly partitioned the data to select and train on 70% (n=255), test on 15% (n=55), and validate on 15% (n=55).

Three, feature engineering, or learning which codon-location variables to include for the most predictive model, was a sequential, sometimes cyclical, culling process to winnow down variables to the smallest set of most highly predictive ones. Many models were built, each with different variables. The most predictive model’s evaluation method for the importance of a codon-location variable was a loss function measured by mean decrease in accuracy (MDA) of variable exclusion. Alternatively, a factor score index combining two load factors – MDA and mean decrease in Gini (MDG) – was considered and rejected because of the MDG metric’s tendency toward overfitting variable importance (Löcher, 2018). The combination of the most effective codon-locations and amino acids was identified through many cycles for the final predictive model. All variables were computed by R, and the top 20 were included for final analysis because variable importance notably decreased after that.

Fourth, the evaluation methods, or test statistics, to compare algorithm performance were (1) area under the receiver operating characteristic curve (AUROC); (2) out-of-bag receiver operating characteristic curve (OOB ROC); and (3) a confusion matrix (i.e., sensitivity, specificity, Type 1 errors, Type 2 errors, and precision). Each test statistic was computed three times for each of the four models. Two tested the sets of 55 data points unseen during algorithmic training. The third test was on the entire dataset. The highest values of the 36 trials (four algorithms x three test statistics x three test datasets) were noted to identify a final predictive model with the highest performance and learn its ingredients.

Fifth, a random forest of classification trees machine-learning algorithm best predicted which coronaviruses could infect humans with or without codon-location substitutions at an average of 98.98% AUROC curve accuracy for the 110 data points the algorithm had never seen before, and 99.63% AUROC for all 365 genomes (including the 255 randomly selected for training). The ordinal ranking of these codon-location variables’ importance, or contribution, to achieve those predictiveness levels identified nine codons that disproportionately impact those outcomes. Those nine codons-locations were then inputs to the machine learning contest described below to learn if a model could be discovered where substitutions of amino acids at those locations could attenuate the virus to be highly improbable to infect humans.

The random-forest algorithm is an ensemble algorithm that averages many differently-sized forests of classification trees ranging from 100 to 10,000 trees to learn the optimum number. Each random forest consisted of a minimum of 3 variables to partition the dataset, which was the square root of the number of input variables. The lowest error rate was at 1000 trees. However, because permutations in variable importance can take longer to stabilize at optimal accuracy, 10000 trees were used to rank variable, or feature, importance. The other three machine-learning models that were run for comparison were: an extreme gradient boost (XGBoost), a support vector machine (SVM) with a nonlinear radial kernel, and deep neural networks with 2 to 13 hidden layers (e.g., 9-2-1 … 9-13-1 networks each with 98 weights).

## Results

Amino-acid substitutions at nine codon-locations attenuated 98.98% to 99.63% (AUROC) of the 365 unique coronavirus genomes studied. The substitutions are alanine to (1) valine at codon 4273; (2) leucine at codon 5077; (3) phenylalanine at codon 2001; (4) leucine at codon 372; (5) proline at codon 354; (6) phenylalanine at codon 2811; (7) phenylalanine at codon 4703; (8) leucine to serine at codon 2333; and (9) threonine to alanine at codon 5131. Further locational analysis found 34% of these changes are in NSP3, 33% in NSP12, 22% in NSP2, and 11% in NSP9. This study suggests codons that encode amino acids in ORF1a, involved in the early control of hosts’ innate and cellular immune response, have a much more critical role in human infectability than those in ORF1b.

## Discussion

### Principal Findings

The principal finding is that by substituting amino acids at nine codon locations, the SARS-CoV-2 virus can be attenuated to not infect humans and protect against 99% of the 365 coronaviruses and variants tested — moreover, amino acids encoded by codons in NSPs 3 and 12 significantly impacted attenuation and human infectability.

There are important insights into the likely functions of these NSPs from prior studies. Generally, the role of NSPs 2-16 is to compose viral replication/transcription complex (RTC). They are targeted to pre-defined subcellular locations. Their interactions with hosts’ cells then determine the replication cycle course.^24 25 26^

NSPs 2-11 appear to provide essential supporting functions enabling the viral RTC process (e.g., cofactors for replication, host immune evasion, modulating intracellular membranes). NSPs 12-16 perform enzymatic functions (e.g., RNA modification, proofreading, and synthesis).^24 27 26^

NSP3, a papain-like proteinase protein, is the largest encoded by coronaviruses (except for the ORF1b polyproteins).^28^ Along with NSP4 and NSP6, it has a transmembrane domain.^29^ NSP3 has been shown to cleave a site between NSP2 and NSP3 and plays a vital protease activity by releasing essential proteins from NSP1, NSP2, and NSP3 for viral activity N-terminal region of ORF1ab.^30^ NSP4 is the second transmembrane domain and interacts with NSP3 and probably host proteins related to membrane re-arrangement.^28^ NSP7 is requisite to form a complex with NSP8 and NSP12 yielding the RNA-polymerase activity of NSP8.^31 28^ NSP8 is a peptide cofactor making a heterodimer with NSP7 (the peptide cofactor), which also complexes with NSP12 to form an RNA-polymerase complex eventually.^32 28^ NSP9 supports the binding of single-stranded RNA.^24^ NSP12 performs RNA synthesis via the RNA-dependent RNA polymerase (RdRP) and two cofactors, NSP7 and NSP8.^24 27 26 32 33^

It also clarifies some evidence in the literature suggesting valine was the most common amino acid driving coronaviruses infecting humans instead plays a more key role in the virulence of the SARS-CoV-2 virus. Researchers believe valine’s (and glycine’s) affinity for calcium leads to the formation of insoluble and rigid calcium oxalate that damages vascular cells, including in the heart. Moreover, they have suggested that amino acids capable of hydrogen bonding may promote weak acids that solubilize this calcium oxalate.^34^

Whether the codons targeted by this machine-learning analysis can be validated in vitro and in vivo remains to be learned and seen; however, are the next essential steps to making a pan-coronavirus globally accessible and halt the coronavirus pandemic. Traditionally, the mechanism for live virus attenuation is codon deoptimization (CDP).^35^ While CDP appears to be efficient and has been used to attenuate viruses for some time, the exact mechanisms that lead to attenuation are unknown. The technique focuses on synthetically recoding viral genomes to destabilize especially mRNA by changing synonymous codons’ positions, increasing suboptimal codon pairs. CDP researchers have noted the need for future research, like here, to determine the functional contribution of individual codon pairs and that such knowledge may lead to safer and more non-reverting attenuated live virus vaccines.^36^

### Comparisons with Prior Work

Prior work regarding coronavirus codons has focused on their amino-acid usage, frequencies, positions, or loci across species at a holistic or broad level. It has also focused on the nucleotide composition of genomes instead of the comparative functionality of codons.^37 38 39 40^ Other research has focused on the statistical properties of amino-acid covariance as possible descriptors or categorizers of viral genomic complexity.^41^ Because of the COVID-19 pandemic, prior work has mainly concentrated on beta coronaviruses (βCoV) instead of the other three genera in the larger *Coronavirdiae* family (e.g., alpha, beta, delta, gamma), like the work here. There has also been substantial research on the functional proteomics of nonstructural proteins (NSPs) in coronaviruses (see Principal Findings); however, little at a codon level of precision. Since 2019, scholarly papers published in academic journals on RNA editing have increased to hundreds per year; however, most are on elements of short-term edits of human RNA for a specific disease or use cases. Recent studies have also explored gene editing of RNA viruses with adenosine deaminases acting on RNA (ADAR) and using machine learning to identify guides for editing RNA viruses using CRISPR-Cas13.^42 21^ Insights gleaned in the present work on codon functionality enabling species-specific infections for many coronaviruses may be more significant strategic value because they are generalizable, informative, and likely actionable in applying basic research to real-world solutions.

### Limitations

While this novel method can precisely locate the codons that enable transmission of emerging viruses to humans at the furthest point upstream at the root amino-acid level, the trinucleotide sequences do not reveal how they work. Therefore, it is a subject area of essential future research to study the structure of these molecules to understand their downstream systems and methods of functioning. Additional future research to evaluate how this method works for other viruses would also be valuable. Understanding the downstream repercussions on proteins of these precise edits, or point mutations, is essential but beyond the scope of this study.

Moreover, no guarantee changing the amino acids at the predicted codon locations would result in a less infective or virulent virus strain. While such changes frequently cause a loss of function, the goal anticipated, sometimes the functional result is minimal or none.^43^

Most of all, like all good moist lab collaborations, it would be invaluable to return these insights to a wet lab to determine in animal models whether editing or substitution of these codons has the effect the machine learning analysis suggests, to introduce loss function in viruses rendering them unlikely or incapable of infecting humans or doing so with significantly less virulence.

## Conclusion

While this study’s insights into better understanding the dynamics and causative codons in human infectability of coronaviruses and how to reverse them to attenuate may be informative, actionable, and have a medically and socially transformative impact, it is the less significant of two conclusions. The more substantial conclusion is proof of new capabilities. Namely, genomic sequences of viruses can be transformed to be analyzed by machine learning algorithms to help explain the phenotypes and functions of codons and proteins. This method, tested and expanded to other pathogens and species, and performed at scale, could yield at least two significant and novel capabilities: (1) collectively across pathogens and species, it could produce the contents of an atlas to serve as a reference for codon or protein phenotypes and functional proteomics (i.e., gain or loss of function blueprints); and, (2) it may serve as a generalizable targeting system for gene editing to create new attenuated virus vaccines (e.g., influenza, HIV, hemorrhagic fevers, etc.) and, similarly, other species too to protect food sources and biodiversity.

## Abbreviations

AUROC: Area under the receiver operating characteristic curve
CART: Classification and regression tree
CDP: Codon deoptimization
CRISPR: Clustered regularly interspaced short palindromic repeats
GDP: Gross domestic product
IBV: Infectious bronchitis virus
OOB: Out-of-bag (e.g., error rate)
ORF: Open reading frame (e.g., gene and polyprotein)
LLM: Linear logistic model
MDA: Mean decrease in accuracy (from variable exclusion in the model)
MDG: Mean decrease in Gini
MERS: Middle East Respiratory Syndrome
NIH US: National Institute of Health
NSP: Nonstructural protein
PDB: Protein databank
RNA: Ribonucleic acid
ROC: Receiver operating characteristic
SVM: Support vector machine
TPA: Third-party annotation

## References

1. Hoecklin, M. (2021, December 1). €93 Billion Spent By Public Sector On COVID Vaccines and Therapeutics in 11 Months, Research Finds. Retrieved from Health Policy Watch: https://healthpolicy-watch.news/81038-2/

2. McKenna, M. (2021, November 1). The Race Is On to Develop a Vaccine Against Every Coronavirus. Wired. Retrieved from https://www.wired.com/story/the-race-is-on-to-develop-a-vaccine-against-every-coronavirus/

3. Center for Systems Science & Engineering (CSSE), Johns Hopkins University (JHU). (2022, February 17). COVID-19 Dashboard. Retrieved from Johns Hopkins University Coronavirus Resource Center: https://coronavirus.jhu.edu/map.html

4. Xie, Y., Xu, E., Bowe, B., Al-Aly, Z. (2022). Long-term cardiovascular outcomes of COVID-19. Nature Medicine. doi:https://doi.org/10.1038/s41591-022-01689-3

5. Piret, J. B. (2021). Pandemics Throughout History. Frontiers in Microbiology, 11. doi:https://doi.org/10.3389/fmicb.2020.631736

6. The IPBES Bureau and Multidisciplinary Expert Panel (MEP). (2020). Workshop Report. IPBES Workshop on Biodiversity and Pandemics (pp. 1–96). Bonn: Intergovernmental Science-Policy Platform on Biodiversity and Ecosystem Services (IPBES).

7. Francis Collins, Director, US National Institute of Health (NIH) (2020, February 4). Influencers: The Dangers of the Coronavirus & How to Stop It. (A. Serwer, Interviewer)

8. Morens, D., Taubenberger, J., Fauci, A. (2022). Universal Coronavirus Vaccines — An Urgent Need. New England Journal of Medicine, 86:297–299. doi:10.1056/NEJMp2118468

9. The Gates Foundation. (2022, January). Bill & Melinda Gates Foundation and Wellcome Pledge US$300 Million to CEPI for COVID-19 Pandemic Response and to Accelerate Epidemic Preparedness. Press Release. Retrieved from https://www.gatesfoundation.org/ideas/media-center/press-releases/2022/01/gates-foundation-wellcome-pledge-300-million-cepi-covid19-pandemic-response

10. Hinshaw, D. (2022, February 19). Bill Gates Says Universal Coronavirus Vaccine Is Possible. The Wall Street Journal. Retrieved from https://www.wsj.com/livecoverage/covid-2021-02-19/card/n5pZK6ZGi0sdT2KoE0rw

11. Luellen, E. (2020). A Machine Learning Explanation of the Pathogen-Immune Relationship of SARS-CoV-2 (COVID-19), and a Model to Predict Immunity and Therapeutic Opportunity: A Comparative Effectiveness Research Study. JMIRx Med, 1(1):e23582. doi:10.2196/23582

12. Batista, C., Hotez, P., Amor, Y., Kim, J., Kaslow, D., Lall, B., & Ergonul, O., et al. (2021). The silent and dangerous inequity around access to COVID-19 testing: A call to action. The Lancet, 43: 101230. doi:https://doi.org/10.1016/j.eclinm.2021.101230

13. Harper, K. (2022, February 10). From obscurity to a Nobel Prize nomination: Houston scientists acclaimed for their patent-free COVID-19 vaccine. The Texas Tribune. Retrieved from https://www.texastribune.org/2022/02/10/corbevax-texas-coronavirus-vaccine/

14. Hafner, M., Yerushalmi, E., Fays, C., Dufresne, E., Van Stolk, C. (2020). COVID-19 and the cost of vaccine nationalism. Santa Monica, CA: RAND.

15. Anthony, S., Johnson, C., Greig, D., Kramer, S.,Che, X., Wells, H., & Hicks, A., et al. (2017). Global patterns in coronavirus diversity. Virus Evolution, 3(1): 1–15.

16. Phillips, J., Jackwood, M., McKinley, E., Thor, S., Hilt, D., Acevedol, N., & Williams, S., et al. (2012). Changes in nonstructural protein 3 are associated with attenuation in avian coronavirus infectious bronchitis virus. Virus Genes, 44(1):63–74. doi:10.1007/s11262-011-0668-7

17. Chan, Y., Teo, T., Utt, A., Tan, J, Amrun, S., Bakar, F., Yee, W., et al. (2019). Mutating chikungunya virus non-structural protein produces potent live-attenuated vaccine candidate. EMBO Molecular Medicine, 11(6):e10092.

18. Li, G., Adam, A., Luo, H., Shan, C., Cao, Z., Fontes-Garfias, C., & Sarathy, V., et al. (2019). An attenuated Zika virus NS4B protein mutant is a potent inducer of antiviral immune responses. NPJ Vaccines, 4(48). doi:https://doi.org/10.1038/s41541-019-0143-3

19. Wang, R., Chen, J., Gao, K., Hozumi, Y., Yin, C., Wei, G. (2021). Analysis of SARS-CoV-2 mutations in the UnitedStates suggests presence of four substrainsand novel variants. Communications Biology, 4(228): 1–14.

20. New York Genome Center. (2020, March 16). New kind of CRISPR technology to target RNA, including RNA viruses like coronavirus. Retrieved from Science Daily: https://www.sciencedaily.com/releases/2020/03/200316141514.htm

21. Wessels, H., Méndez-Mancilla, A., Guo, X., Legut, M., Daniloski, Z., Sanjana, N. (2020). Massively parallel Cas13 screens reveal principles for guide RNA design. Nature Biotechnology, 38: 722–727.

22. de Breyne, S., Vindry, C., Guillin, O., Condé, L., Mure, F., Gruffat, H. & Chavatte, L., et al. (2020). Translational control of coronaviruses. Nucleic Acids Research, 48(22): 12502–12522.

23. NCBI. (2021, April 21). Nucleotide. Retrieved from NCBI: https://www.ncbi.nlm.nih.gov/nucleotide/

24. V’kovski, P., Kratzel, A., Steiner, S., Stalder, H., Thiel, V. (2021). Coronavirus biology and replication: implications for SARS-CoV-2. Nature Reviews Microbiology, 19: 155–170.

25. Sims, A., Ostermann, J., Denison, M. (2000). Mouse hepatitis virus replicase proteins associate with two distinct populations of intracellular membranes. Journal of Virology, 74: 5647–5654.

26. Snijder, E., Decroly, E., Ziebuhr, J. (2016). The nonstructural proteins directing coronavirus RNA synthesis and processing. Advances in Virus Research, 96: 59–126.

27. Perlman, S., Netland, J. (2009). Coronaviruses post-SARS: update on replication and pathogenesis. Nature Reviews Microbiology, 7: 439–450.

28. Yoshimoto, F. (2020). The proteins of severe acute respiratory syndrome coronavirus-2 (SARS CoV-2 or n-COV19), the cause of COVID-19. The Protein Journal, 39: 198–216.

29. Sakai, Y., Kawachi, K., Terada, Y., Omori, H., Matsuura, Y., Kamitani, W. (2017). Two-amino acids change in the nsp4 of SARS coronavirus abolishes viral replication. Virology, 510: 165–174.

30. Lei, J., Kusov, Y., Hilgenfeld, R. (2018). Nsp3 of coronaviruses: structures and functions of a large multi-domain protein. Antirivrual Research, 149: 58–74.

31. te Velthuis, A., van de Worm, S., Snijder, E. (2012). The SARS-coronavirus nsp7+nsp8 complex is a unique multimeric RNA polymerase capable of both de novo initiation and primer extension. Nucleic Acids Research, 40: 1737–1747.

32. Gao, Y., Yan, L., Huang, Y., Liu, F., Zhao, Y., Cao, L., & Wang, T., et al. (2020). Structure of the RNA-dependent RNA polymerase from COVID-19 virus. Science, 368(6492): 779–782.

33. Tvarogová, J., Madhugiri, R., Bylapudi, G., Ferguson, L., Karl, N., Ziebuhr, J. (2019). Identification and characterization of a human coronavirus 229E Nonstructural protein 8-associated RNA 3′-terminal adenylyltransferase activity. Journal of Virology, 93(12). Retrieved from https://doi.org/10.1128/JVI.00291-19

34. Lu, X., Ran, H., Yang, Y., Liu, Q. (2020, May 7). A highly conserved cryptic epitope in the receptor binding domains of SARS-CoV-2 and SARS-CoV. Retrieved from Science: https://science.sciencemag.org/content/368/6491/630/tab-e-letters

35. Vignuzzi, M., Wendt, E., Andino, R. (2008). Engineering attenuated virus vaccines by controlling replication fidelity. Nature Medicine, 14: 154–161.

36. Groenke, N., Trimpert, J., Merz, S., Conradie, A., Wyler, E., Zhang, H., & Hazapis, O., et al. (2020). Mechanism of virus attenuation by codon pair deoptimization. Cell Reports, 31(4): 107586.

37. Tort, F., Castells, M., Cristina, J. (2020). A comprehensive analysis of genome composition and codon usage patterns of emerging coronaviruses. Virus Research, 283: 197976.

38. Hussain, S., Shinu, P., Islam, M., Chohan, M., Rasool, S. (2020). Analysis of codon usage and nucleotide bias in Middle East Respiratory Syndrome coronavirus genes. Evolutionary Bioinformatics, 16: 1–13.

39. Khandia, R., Singhal, S., Kumar, U., Ansari, A., Tiwari, R., Dhama, K., & Das, J., et al. (2019). Analysis of Nipha virus codon usage and adaption to hosts. Frontiers in Microbiology, 10: 886.

40. Pandit, A., Sinha, S. (2011). Differential trends in the codon usage patterns in HIV-1 genes. PLoS ONE, 6(12): e28889.

41. Sruthi, C., Prakash, M. (2019). Statistical characteristics of amino acid covariance as possible descriptors of viral genomic complexity. Nature, 9: 18410.

42. Reardon, S. (2020, February 4). News feature: Step aside CRISPR, RNA editing is taking off. Nature. Retrieved from https://www.nature.com/articles/d41586-020-00272-5

43. National Human Genome Research Institute. (2021, June 1). Missense Mutation. Retrieved from National Human Genome Research Institute: https://www.genome.gov/genetics-glossary/Missense-Mutation

